# Excess mortality in Guadeloupe and Martinique, islands of the French West Indies, during the chikungunya epidemic of 2014

**DOI:** 10.1101/228445

**Authors:** A. R. R. Freitas, P. M. Alarcon-Elbal, M. R. Donalisio

**Affiliations:** Faculdade de Medicina São Leopoldo Mandic, Campinas, Sao Paulo, Brazil.; Universidad Iberoamericana, Instituto de Medicina Tropical & Salud Global, Santo Domingo, Dominican Republic.; State University of Campinas, Faculty of Medical Sciences, Public Health, Campinas, São Paulo.

## Abstract

In some chikugunya epidemics, deaths are not fully captured by the traditional surveillance system, based on case reports and death reports. This is a time series study to evaluate the excess of mortality associated with epidemic of chikungunya virus (CHIKV) in Guadeloupe and Martinique, Antilles, 2014. The population (total 784,097 inhabitants) and mortality data estimated by sex and age were accessed at the Institut National de la Statistique et des Etudes Economiques - France. Age adjusted mortality rates were calculated also in Reunion, Indian Ocean for comparison. Epidemiological data on CHIKV (cases, hospitalizations, and deaths) were obtained in the official epidemiological reports of the Cellule de Institut de Veille Sanitaire - France. The excess of deaths for each month in 2014 and 2015 was the difference between the expected and observed deaths for all age groups, considering the 99% confidence interval threshold. Pearson coefficient of correlation between monthly excess of deaths and reported cases of chikungunya show a strong correlation (R = 0.81, p <0.005), also with a 1-month lag (R = 0.87, p <0.001), and between monthly rates of hospitalization for CHIKV and the excess of deaths with a delay of 1 month (R = 0.87, p <0.0005).The peak of the epidemic occurred in the month with the highest mortality, returning to normal soon after the end of the CHIKV epidemic. The overall mortality estimated by this method (639 deaths) was about 4 times greater than that obtained through death declarations (160 deaths). Excess mortality increased with age. Although etiological diagnosis of all deaths associated with CHIKV infection is not possible, already well-known statistical tools can contribute to an evaluation of the impact of this virus on the mortality and morbidity in the different age groups.

## Introduction

The mosquito-borne chikungunya virus (CHIKV) affects human health around the world, being the main vectors *Aedes aegypti* and *Aedes albopictus*. From its discovery in 1953 in Uganda to the first years of the 21st century CHIKV was recognized as an arbovirus causing small outbreaks in Asia and Africa, with high attack rates but with a low case fatality rate. After 2005, there has been a significant increase in the number of cases and an expansion of the transmission area in several regions of the world, including continental Europe [1,2]. In 2005 and 2006 a major epidemic hit the Reunion Island, a French overseas department located in the Indian Ocean, when severe cases and deaths were described [3,4], In 2006 266,000 symptomatic patients and 255 deaths were estimated, with a fatality rate of 1 death for every 1,000 cases. Since then, several fatal cases have been described in the literature[5–9].

At the end of 2013 an autochthonous transmission of CHIKV was identified in Saint Martin, Guadeloupe and Martinique [10]. The transmission quickly spread through almost all the islands of the Caribbean and also to continental America. By the end of the 2014 epidemic, 1,118,578 CHIKV cases were reported in the Americas; a total of 194 deaths were informed to PAHO, which results in a lethality rate of 0.02%, much lower than reported in the literature [11]. *Aedes aegypti* was the suspected vector of the arbovirus since there is no *Ae.albopictus* on these islands. In fact, a recent study has shown for the first time that CHIKV infections occurred in natural populations of *Ae. aegypti* on Martinique [12].

By the end of the epidemics in Guadeloupe and Martinique a total of 153,400 cases were reported, for an estimated of 306,800 CHIKV symptomatic cases [13]. Considering these data, the fatality rate in Guadeloupe and Martinique would be respectively 0.04% and 0.06%, practically half of what was observed in Reunion in 2006. In view of the fact that the proportion of elderly population of Reunion Island in 2006 was 9.6% and in Guadeloupe, 15.7% and in Martinique, 17.6%; [14], and also that CHIKV deaths occur at more advanced ages, it would be expected a higher mortality in Martinique and Guadeloupe when compared with Reunion.

In Reunion Island, the excess deaths during the CHIKV epidemic was 267 deceases [15], very close to 255 that was found in the death certificates. Differently from Brazil, Ahmedabad, Andaman and Nicobar Islands (belonging to India) and Mauritius where a significant excess of deaths that did not coincide with deaths reported for epidemiological surveillance system [16–20]. Thus, some inconsistent data on estimated mortality in CHIK circulation regions have been reported.

The objective of this study was to assess the excess mortality by sex and age group in the French overseas departments of the Caribbean, Guadeloupe and Martinique, during the 2014 epidemic year of CHIKV using the official data from the Institut National de La Statistique et des Etudes Economiques (INSEE). We also compare results with official data on the 2005-2006 epidemic on Reunion Island. Our hypothesis is that there may have been an excess of deaths above that identified by death certificates.

## Methods

A time series study of reported incidence rates of CHIKV and mortality by sex and age group was conducted in Guadeloupe and Martinique, in the period between 2011 and 2015.

## Locality

Guadeloupe and Martinique are French overseas departments in the Caribbean, are set of islands of tropical climate (Af, according with Köppen Climate Classification System) located in the Lesser Antilles of the French West Indies, with populations of 400,186 and 383,911 inhabitants in 2014, respectively [14].We analyzed the two French overseas departments together because they had a similar climate, epidemiological and sociodemographic profile, and the CHIKV epidemic was practically simultaneous in both territories [13].

## Data collect

The population data estimated by sex and age group, and mortality data were accessed at the INSEE of the French government [21]. Epidemiological data on CHIKV as number of cases, hospitalizations and deaths were obtained in the official epidemiological reports of the Cellule de Institut de Veille Sanitaire (l’InVS) en Region of the Government of France [13]. Mortality data during the 2006 epidemic in Reunion were retrieved through literature [8] based on official surveillance [3]. As the influenza virus is known to be a cause of seasonal increase in overall mortality, we also collect monthly data from sentinel surveillance of influenza-like syndrome in the bulletins de l’InVS [22,23]. Numerical data were extracted from the graphs of bulletins using the program Engauge Digitizer [24].

## Statistical analysis

The monthly number of deaths expected for 2014 and 2015 by age group and sex was calculated by applying the average mortality rates per month of the reference period for the estimated population of the French departments of Guadeloupe and Martinique in 2014 and 2015. The reference period used was three years prior to 2013. The excess of deaths for each month in 2014 and 2015 was calculated as the difference between the monthly expected deaths for all age groups and the total deaths observed per month. The conservative confidence interval of 99% of the monthly number of deaths was also calculated. The excess of obits were also calculated for the whole years 2014 and for each age group and sex. The excess age-standardized death rates (ASMR) were calculated for the year 2014 by the direct method using the formula below (formula 1) using the WHO standard population and the excess of deaths by age group of 2014 [25].

Formula 1:

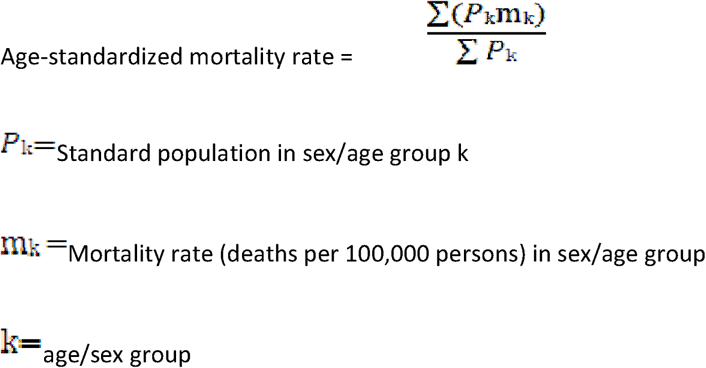

The mortality rate by age group for the whole year 2006 of Reunion Island was estimated by extrapolating the mortality rate observed in the period from January to April [3] for deaths observed throughout the year 2006 [8] assuming the same proportion of deaths by age group. Statistical analyzes were performed using the IBM^®^ SPSS^®^ 24.0 software.

## Results

### CHIKV epidemic curve and monthly mortality

The monthly reported cases of CHIKV, the number of hospitalizations and excess deaths in 2014 in the departments of Guadeloupe and Martinique diagnosed by sentinel doctors is presented in Table 1. We also show the estimates made by InVS of clinical cases of CHIKV, in Guadeloupe and Martinique in the year 2014. There was a strong correlation between the monthly incidence rates of CHIKV and the excess of deaths (R = 0.81, p <0.005), and ever higher with a 1-month lag between the CHIKV cases and the excess of deaths (R = 0.87, p <0.001, Table1). We observed a strong correlation between the monthly rates of hospitalization for CHIKV and the excess of deaths with a delay of 1 month (R = 0.87, p <0.0005) while a moderate correlation was found between the monthly hospitalization rates and the excess of deaths (R = 0.66, p <0.05) (Table1)‥

**Table 1.** Monthly cases of chikungunya, influenza like illness and excess deaths (Guadaloupe and Martinique, 2014)

|  |  |  | Chikungunya<br>Cases<br>(clinically<br>diagnosed -<br>sentinel<br>physicians) | Total of<br>Estimated<br>Chikungnuya<br>Clinical<br>Cases | Hospitalization | Influenza Like<br>Illness<br>(sentinel<br>surveillance) | Excess<br>deaths |
| --- | --- | --- | --- | --- | --- | --- | --- |
| jan |  |  | 2,161 | 4,322 | 65 | 5,608 | 5 |
| fev |  |  | 4,102 | 8,204 | 101 | 5,475 | -12 |
| mar |  |  | 7,924 | 15,848 | 138 | 5,136 | 13 |
| abr |  |  | 18,254 | 36,508 | 258 | 3,260 | <u>40</u> |
| mai |  |  | 27,471 | 54,942 | 323 | 897 | <u>61</u> |
| jun |  |  | 37,936 | 75,871 | 315 | 816 | <u>159</u> |
| jul |  |  | 26,197 | 52,395 | 260 | 610 | <u>105</u> |
| ago |  |  | 12,716 | 25,432 | 198 | 570 | <u>105</u> |
| set |  |  | 7,891 | 15,782 | 125 | 743 | <u>49</u> |
| out |  |  | 5,871 | 11,742 | 59 | 1,983 | <u>47</u> |
| nov |  |  | 2,326 | 4,652 | 37 | 1,857 | <u>36</u> |
| dez |  |  | 851 | 1,702 | 10 | 1,547 | 31 |
| Total |  |  | 153,700 | 307,400 | 1,888 |  | 639 |
| Correlati<br>on<br>(Pearson<br>) with<br>monthly<br>excess of<br>death | without<br>lag | R |  | 0.806 | 0.664 | -0.76* |  |
|  |  | *p* |  | <0.005 | <0.05 | <0.005 |  |
|  | 1 month<br>lag | *R* |  | 0.868 | 0.872 | -0.66* |  |
|  |  | *p* |  | <0.001 | <0.0005 | <0.05 |  |
|  | 2 month<br>lag | R |  | 0.639 | 0.763 | -0.197 |  |
|  |  | *p* |  | <0.05 | <0.05 | 0.586 |  |
| <u>Months with number of death above upper 99% confidence interval</u> |  |  |  |  |  |  |  |

\* Without biological plausibility

Figure 1 shows the number of deaths expected and observed per month, above the upper limit of the 99% confidence interval in Guadeloupe and Martinique. We also present the 99% confidence interval and the number of monthly cases of CHIKV estimated by InVS. The peak of the epidemic was in June, as was the month in which the highest number of deaths occurred. Deaths remained above the 99% confidence interval in the months of April to November period of highest incidence of CHIKV. In the months with a small number of cases of CHIKV in 2014 (January to March and December) and in 2015 the number of monthly deaths per CHIKV was below the upper limit of the 99% confidence interval.

**Figure 1.**
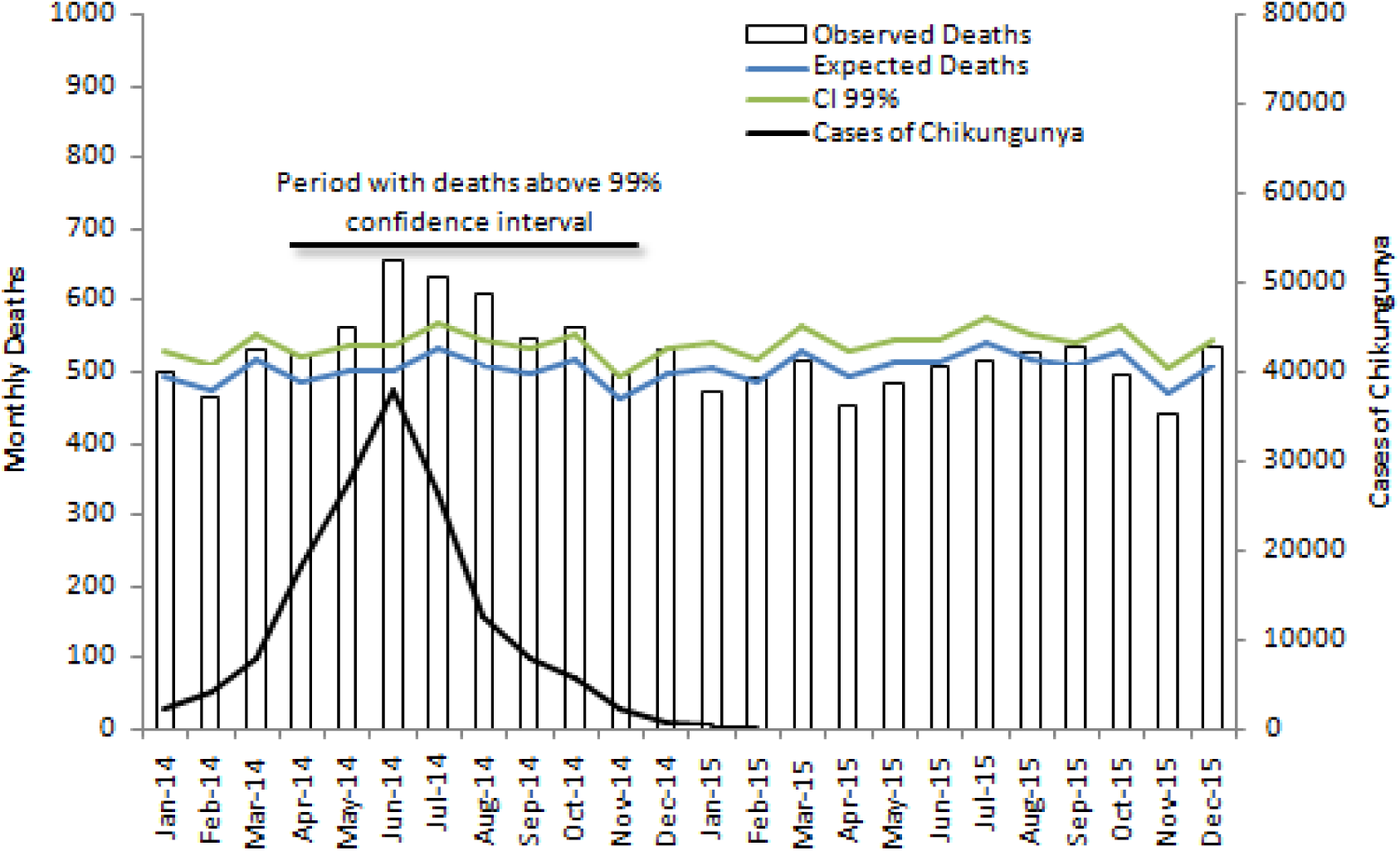
Cases of chikungunya, expected, observed monthly deaths and upper limit 99% confidence interval (Guadaloupe and Martinique 2014-2015)

### Mortality by sex and age group

Except for the under-19 age group, all others had a higher number of deaths than expected. ASMR was above the upper limit of the 99% confidence interval in all age groups over 20 years of age, with the exception of the age range between 40 and 59 years (Table 2); the same pattern was observed in both sexes (Figure 2). the excess mortality The excess of deaths increased with age in both sexes. The values found for the excess ASMR in Guadeloupe and Martinique were not very different from the ASMR CHIKV observed in Reunion Island in 2006, estimated using death certificates (Table 2). In the year 2015, ASMR were lower than observed in 2014 and were below the upper limit of the confidence interval in all age groups.

**Table 2.**
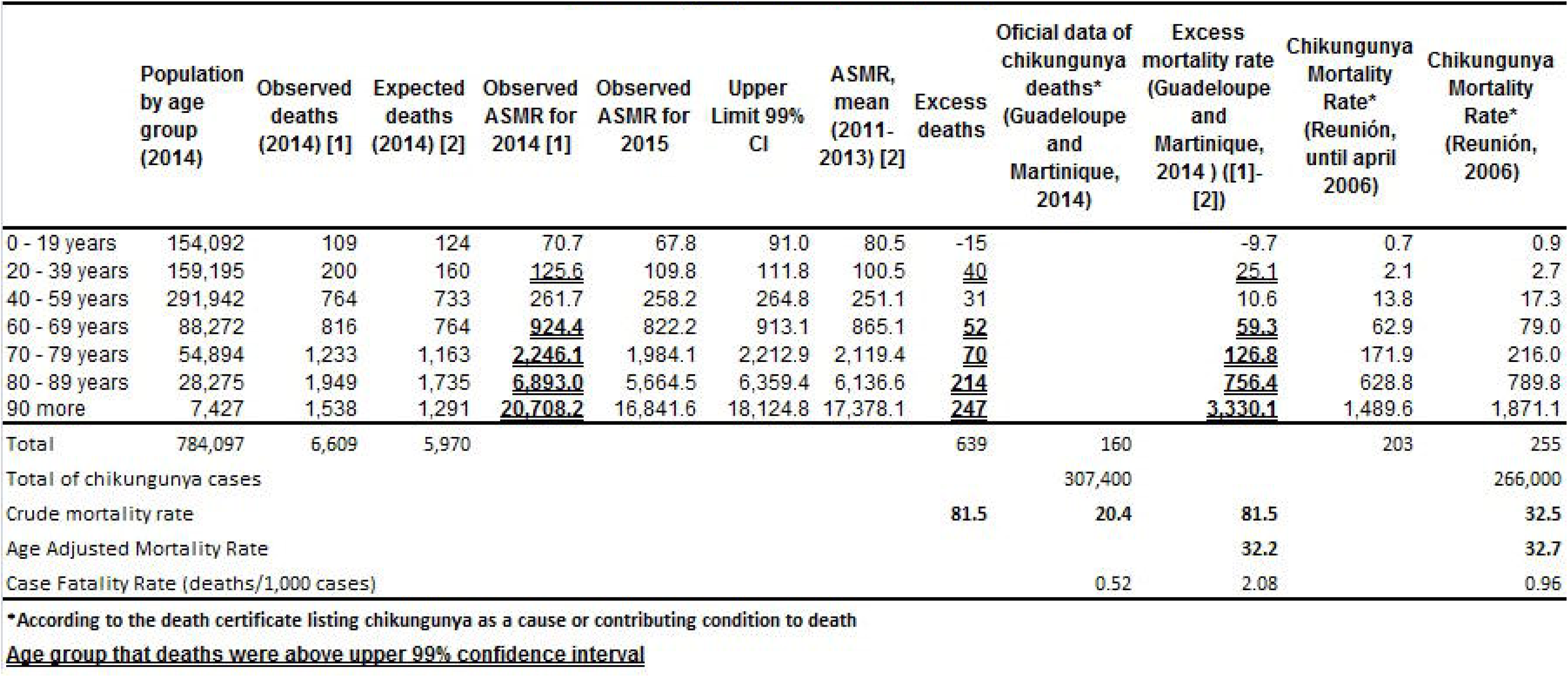
Mortality data, observed and estimated in Guadaloupe and Martinique (2014) and observed in Reunion Islands (2006). (ASMR: Age-Specific Mortality Rate)

**Figure 2.**
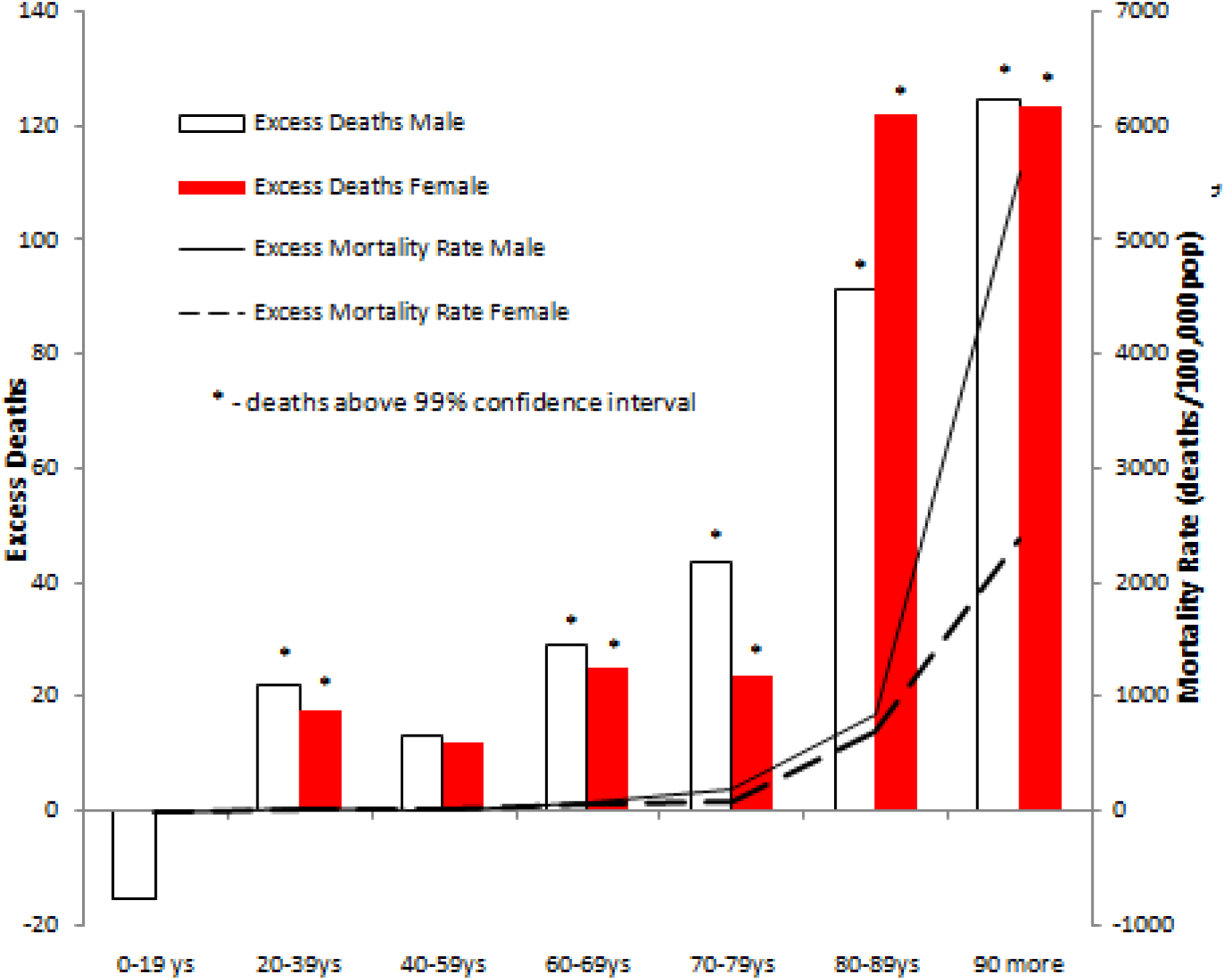
Excess deaths and mortality rate by sex and age group (Guadaloupe and Martinique, 2014)

### Excess mortality, age-adjusted mortality rate and fatality rate

During 2014 in Guadeloupe and Martinique there was 639 deaths more than expected resulting in an excess mortality rate of 81.5 deaths per 100,000 inhabitants, 4 times greater than that reported to InVS through death certificates, and 2.5 times greater than observed in Reunion in 2006. The excess mortality adjusted by WHO standard age (world average 2000-2025) was 32.2 deaths per 100,000 population (Table 2) very close to that observed in Reunionin 2006 (32.7 deaths per 100,000 population). The fatality rate in 2014 was 2.08 deaths per 1,000 cases of CHIKV (Table 2), practically twice that registered in 2006 in Reunion (0.96 deaths per 1,000 cases of CHIKV). These differences are probably due to the fact that the proportion of older people in the French Antilles in 2014 was higher than in the Reunion Island in 2006.

## Discussion

The time series analysis of deaths in the Guadeloupe and Martinique departments shows that the standardized mortality rate increased above the 99% confidence interval during the CHIKV epidemic of 2014, rates returning to normal soon after the end of the epidemic. The peak of deaths occurred in June, simultaneous with the peak of CHIKV cases. There was a strong temporal correlation between the occurrence of excess deaths, cases of CHIKV and hospitalization. This correlation was stronger when we used 1 month of delay between deaths for the occurrence of clinical cases and hospitalizations. This lag between the occurrence of the clinically identified cases and the deaths is compatible with long periods of hospitalization observed as consequence of clinical complications, especially among the elderly with prevalent chronic diseases [26]. Correlation between monthly hospitalization rates and the excess of deaths reinforces this hypothesis. Excess mortality increased with age, but there were excess deaths in the age groups of 20 to 39 years, being above the upper limit of the 99% confidence interval. This same mortality pattern was observed in studies that analyzed the CHIKV cause or cause of death as reported in the certificates [3] and in studies that used laboratory test results (RT-PCR, viral isolation or IgM) to confirm the disease [4,7],The excess mortality rate by age group in the Guadeloupe and Martinique in 2014 (Table 2) was very close to the specific mortality by CHIKV observed in Reunion in 2006, considering the declarations of deaths [3,8]. These findings reinforce our hypothesis that increased mortality was a consequence of the CHIKV epidemic.

There was no positive correlation between the increase in deaths and circulation of influenza virus, another frequent cause of increase in overall mortality. There were no other phenomena in these islands that could be related to an increase in mortality in this period. An epidemic of dengue that began in mid-2013 ended on March 2, 2014 in Guadeloupe and on April 20, 2014 in Martinique [27], and apparently there was no significant circulation of other arboviruses.

A total of 639 excess deaths were associated with this CHIKV epidemic, 4 times higher than the number of death certificates that mentioned CHIKV as cause. In the 2006 Reunion epidemic the estimated excess was very similar to the number of deaths in which there was mention of CHIKV in the death certificates as the basic cause or contributing cause [8,15]. Perhaps the fact that it was the first major epidemic in French territory, the etiologic investigation and the detailing in the preparation of the certificate of obit have been made more carefully by the health professionals. Recent studies carried out in the Antilles have shown that the elderly present an acute clinical picture different from the younger ones and this may help to explain, at least partially, the difficulty in the etiological diagnosis of the deaths in elder during the CHIKV lack of knowledge of the severe forms of CHIKV that may present complications of a very diverse nature such as myocarditis, arrhythmias, hepatitis and encephalitis [4,28]. Another reason that may hamper the attribution of death to CHIKV is the fact that many patients may die as a consequence of exacerbation of underlying diseases, such as cardiovascular diseases [4], Although some researchers still considers that the deaths caused by CHIKV are rare [29], data from Martinique and Guadalupe departments suggest the opposite. This lack of consensus on the ability of the CHIKV lead to death may conduct doctors not to attribute a death to the CHIKV causing this underreporting as observed in our study.

Considering that the InVS estimated the number of cases of symptomatic patients at 307,400, the estimated lethality was 2 deaths per 1,000 cases, 10 times higher than that reported by all the countries that make up the PAHO in the year 2014 [11],In 2006 the fatality rate attributed to CHIKV in the Reunion Island was 1 deaths for 1,000 CHIKV cases, half of this rate was observed in the Antilles in 2014. This divergence probably was due to the difference in the age profile since the proportion of elderly people in the Antilles in 2014 was higher than in the Reunion in 2006 [8]. Published study evaluating excess mortality in India in 2006 raised the hypothesis that deaths could be linked to increased virulence related to mutations in the East/Central /South African (ECSA) lineage of the CHIKV [17]. Our findings suggest that the Antillean epidemic in 2014, caused by a CHIKV of the Asian lineage, had a mortality by age group very close to that observed in the 2006 Reunion epidemic, that were caused by the ECSA lineage.

The main limitation of this study is associated with the design that is based on a temporal series of deaths, with no etiological definition. Other phenomena may have been responsible for this increase in overall mortality in Guadeloupe and Martinique. However, the temporal pattern and age of the deaths are very similar to those found in epidemics that occurred in other localities in which laboratory criteria were used to confirm cases. Our results are compatible with the association of CHIKV transmission with excess of mortality with higher impact in the elderly. The lack of etiological diagnosis of the infection may not mean that the disease does not increase risk of death. A situation of this nature occurs with influenza in which only a small part of the deaths is diagnosed as such. Most influenza-related deaths result from secondary bacterial pneumonia or from decompensation of underlying diseases caused by viral infection and death is not classified as due to influenza, thus, the impact of influenza on mortality is estimated by assessing the excess of influenza mortality [30,31]. Using similar criteria of present work the excess of deaths caused by influenza among those over 65 years is 73 deaths per 100,000 per year [30], with a simple calculation with the data of the present study we conclude that the excess of deaths associated with a CHIKV epidemic is 324 per 100,000 inhabitants over 60 years or 4.4 times higher than the annual average of mortality associated with influenza[30].

Mortality and fatality are very expressive indicators for assessing disease burden and important for use in prioritizing public health, including vaccine production. Official documents have placed CHIK as a disease less lethal than dengue, stating that deaths are rare [32], Studies in Reunion Island and Ahmedabad have shown that a significant proportion of deaths occurred in patients with relatively common and non-severe conditions such as high blood pressure and diabetes, and even without underlying diseases [4,7],

Between 2004 and 2015, in Guadeloupe and Martinique 49 deaths was attributed to dengue and 17,1550 cases was estimated resulting in a fatality rate of 0.29 deaths per 1,000 cases [27], According to official data from the InVS, that considers death certificates, the number of deaths per CHIKV in Guadeloupe and Martinique in a single year is 3.3 times higher (160 deaths) than the accumulated dengue in 10 years and fatality rate almost twice as high (0.52 deaths per 1,000 cases). Using the excess mortality criteria, the mortality associated with the CHIKV epidemic in a single year would be 13 times greater than that accumulated in 10 years of dengue epidemics in the same locality, and the fatality rate would be 7.7 times greater than dengue. Although it is not possible to make the etiological diagnosis of all the cases of deaths associated with CHIKV infection, already well-known statistical tools can contribute to an evaluation of the impact of this virus on the mortality in the different age groups, as in the deaths caused by extreme weather phenomena, seasonal and pandemic influenza[31,33,34].

Knowledge of the actual potential for morbidity and mortality of an epidemic viral infection such as CHIKV and the most vulnerable groups may help health professionals during epidemics.

